# Rapid identification of key genes for the rod-shaped morphology in bacteria using multi-species genomes

**DOI:** 10.1101/2023.07.14.548972

**Authors:** Qi Liu, Haida Liu, Chuangchuang Xu, Jianqiang Shi, Yanghe Xie, Shunli Hu, Guomin Han

**Author notes:** Correspondence (G.H.).

## Abstract

Accurate identification of key genes is pivotal in biological research. Here, we introduce machine learning to the field of functional gene identification, enabling precise prediction of bacterial shape based on genomic information. Our machine learning model successfully predicts bacterial shape, and we determine the influence of various protein domains on shape using the model. This approach facilitates the identification of candidate genes involved in regulating bacterial shape. Through targeted knockout experiments on eight potential key regulatory genes (*pal, yicC, mreB, tolQ, ftsX, amiC, yddB*, and *rpoZ*) in *Escherichia coli*, we observe significant alterations in rod-shaped morphology upon individual knockout of *pal* and *mreB* genes. *E. coli* transitions from rod-shaped to spherical or cell wall-deficient protoplasmic states. Experimental validations validate the robustness of our newly developed method. This study establishes an innovative avenue for exploring functional genes, harnessing large-scale genomic information to promptly uncover key genes governing shared traits across species.

## Introduction

Identification of key genes plays a pivotal role in biological research as genes are fundamental molecules that govern diverse life processes and traits[1-3]. Unraveling their functions is crucial for elucidating the basic principles of life, investigating biological development and evolution, and uncovering the underlying mechanisms of disease occurrence[2, 4]. With advancements in biological technology, several commonly used methods have been developed to investigate gene functions, including mutant screening, map-based cloning, and genome-wide association analysis (GWAS)[5-8]. Through these methods, numerous important genes have been identified, including those responsible for determining bacterial shapes[9].

Bacteria exhibit diverse morphological types, including spherical, rod-shaped, spiral, stalked, square, star-shaped bacteria, and more[10, 11]. However, the primary categories are spherical, rod-shaped, and spiral bacteria[11]. The selection of bacterial shape provides distinct advantages for survival as it influences cell motility, attachment, colonization in new environments, and defense against predation[12]. The shape of bacteria plays a significant role in determining their adaptive strategies and overall fitness within their ecological niches[13].

The mechanisms of bacterial morphogenesis are intricate and involve regulation at multiple levels. Recent studies have revealed that the maintenance and alteration of bacterial shape are influenced by factors such as the cell wall, environmental conditions, and gene regulation[14-16]. Many genes have been identified to encode enzymes and proteins that play a role in regulating bacterial morphogenesis, controlling cell wall growth, and shaping [9, 14, 15, 17]. In most bacteria, the rod-like morphology is maintained by the elongasome, which consists of essential proteins including PBP2, MreBCD, RodA, and others[17]. These proteins mediate peptidoglycan synthesis and insertion into the cylindrical part of the cell wall[17]. Notably, penicillin-binding protein (PBP) is crucial for bacterial cell wall synthesis and repair. It not only participates in the formation of peptidoglycan chains but also cross-links these chains, influencing the composition, structure, and shape of the bacterial cell wall[17]. The *MreB* and *MreC* genes encode proteins that play significant roles in maintaining bacterial shape[9, 14]. MreB is a cytoskeletal protein belonging to the actin superfamily, along with microtubulin and cytoskeletal proteins. Its spatial arrangement is associated with cell shape and is involved in cell wall synthesis and cytoplasmic distribution during cell division[14]. For example, in *Escherichia coli*, a rod-shaped bacterium, MreB is localized in a curvature-dependent manner and aids in maintaining cell shape by coordinating cell wall insertion [15]. Mutations in the *MreB*(A125V) and *MreB*(A174T) genes lead to variations in cell width, with the mutant cells being thinner or thicker than wild-type cells. The rotation speed of MreB also affects cell width, with faster rotation resulting in narrower cells [18]. MreC is a membrane-bound protein that interacts with the cell membrane and works synergistically with MreB to maintain cell shape. Deletion of the *MreC* gene leads to altered cell shape and abnormal cell wall synthesis[19]. Other proteins such as *FtsZ*-encoded proteins[20, 21], *RodZ*-encoded proteins[22], and Bactofilins are also involved in bacterial cell division and shape maintenance[23]. RodZ proteins regulate the spatial structure of MreB by interacting with other proteins, helping to maintain rod-like cell shape. Bactofilins, small β-helix proteins, form cytoskeletal filaments and play roles in *Caulobacter crescentus* and *Myxococcus xanthus* in chromosome segregation and movement[20, 22, 23].

The current methods used to identify key functional genes are primarily based on the analysis of individual species. While significant advancements have been made, these methods have limitations due to various factors. One limitation arises from the random distribution of mutations generated by chemical or physical mutagenesis in many organisms[7, 8]. Some mutations may be ineffective or unrelated to the specific study, necessitating considerable time and effort to identify and screen for meaningful mutants and subsequently mine them to uncover key genes[7]. Additionally, generating and maintaining mutant libraries require substantial resources and time, as well as the challenge of dealing with lethal mutations and the inability to cover all gene mutations comprehensively. Map-based cloning has emerged as another effective method for identifying functional genes in recent decades[5, 24, 25], but it often requires several years to obtain desirable results. With the rapid development of genome sequencing technology, GWAS has become a functional gene identification method based on population genetics[6, 26-28]. However, conducting GWAS necessitates substantial financial resources, effort, and time to collect a sufficient amount of materials, such as different strains of the same species or various plant and animal varieties[26-29]. Only then can excellent genotype be identified through GWAS[6]. It is important to note that existing methods for identifying key functional genes have mainly focused on analyzing single species. Despite significant progress, many genes have multifunctional properties, and there are still numerous bacteria, fungi, plants, and animals where the function of certain genes predicted from genomics remains entirely unknown. Additionally, a gene with a known function may also possess other unknown functions. This highlights the ongoing challenge of comprehensively understanding gene functions across diverse organisms.

In this study, we developed a method to identify key genes that regulate bacterial rod shape using information extracted from a large number of bacterial genomes. We established a comprehensive approach that combines bioinformatics analysis and experimental techniques to identify and validate the candidate genes involved in controlling bacterial rod shape.

## Results

### Construction of machine learning model

We employed a dataset consisting of 4,847 bacteria with matched genomic and trait information to train and test our model. However, the initial results were not satisfactory, as indicated by the low Kappa coefficient and a support vector machine accuracy of 0.59, which represents moderate prediction accuracy (Supplementary Figure 1). To address this issue, we adjusted the number of bacteria in different proportions and ultimately utilized a total of 3,750 bacteria from the available dataset (Table 1). For the training set, we included 854 cocci, 999 bacilli, 589 spirochetes, and 373 other bacteria. The test set consisted of 284 cocci, 331 bacilli, 196 spirochetes, and 124 other bacteria. We applied five machine learning algorithms, namely the support vector machine algorithm, conditional inference tree algorithm, decision tree algorithm, Naive Bayes algorithm, and random forest algorithm, to train the model using the aforementioned training set.

**Table 1.**
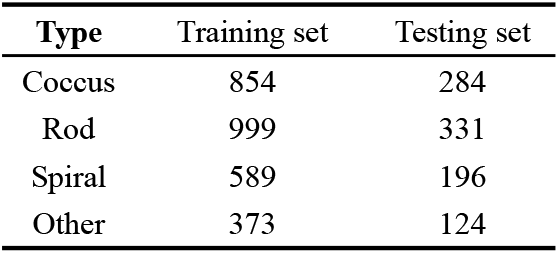
Optimal model grouping list.

**Figure 1.**
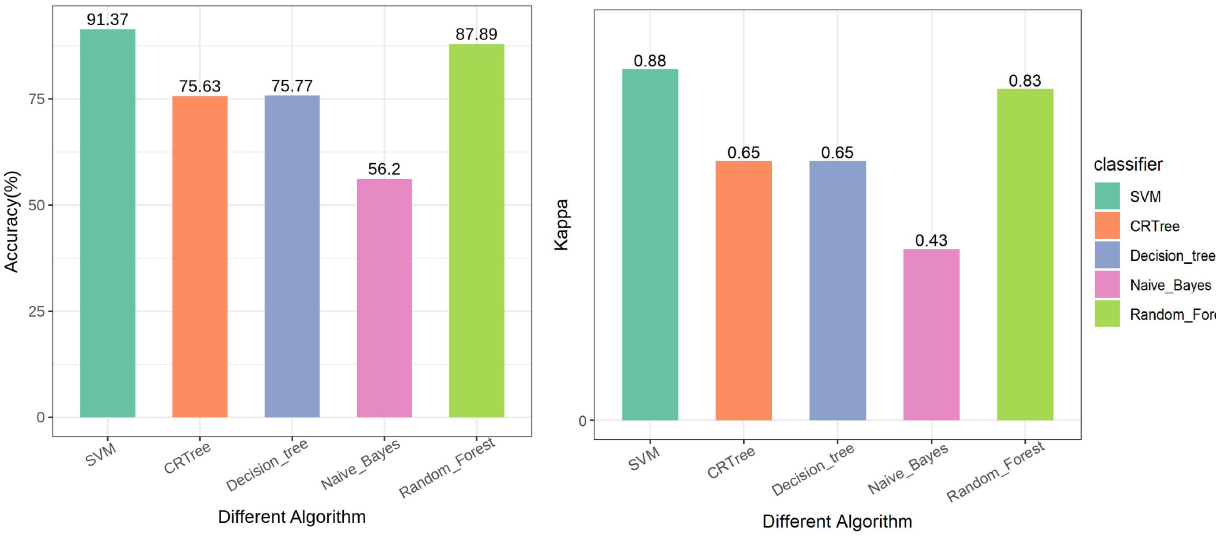
Predicting results of five machine learning algorithms on the training set Note: SVM: Support vector machine algorithm, CTree: conditional inference tree algorithm, Decision_tree: decision tree algorithm, Naive_Bayes: Naive Bayes algorithm, Random_Forest: random forest algorithm.

The overall performance of the optimized model was satisfactory, particularly with the support vector machine and random forest algorithms. The accuracy of model training reached 91.37% and 87.89%, respectively, with both algorithms yielding kappa coefficients above 0.81 (Figure 1). The recall rates for cocci, bacilli, spirochetes, and other bacteria also demonstrated good performance. When predicting the test set data using the trained model, the support vector machine algorithm achieved a prediction accuracy of 86.31%, while the random forest algorithm achieved an accuracy of 94.76% (Figure 2). The random forest algorithm outperformed the support vector machine algorithm in the prediction of the test set. Notably, the recall rates for coccus, bacillus, spirochete, and other bacteria were 97.18%, 92.75%, 98.98%, and 87.90%, respectively, which were substantially higher than the other four algorithms (Figure 3). The high kappa coefficient of 0.93 indicated that the classification results of the random forest algorithm closely matched the actual data (Figure 2&Figure 3).

**Figure 2.**
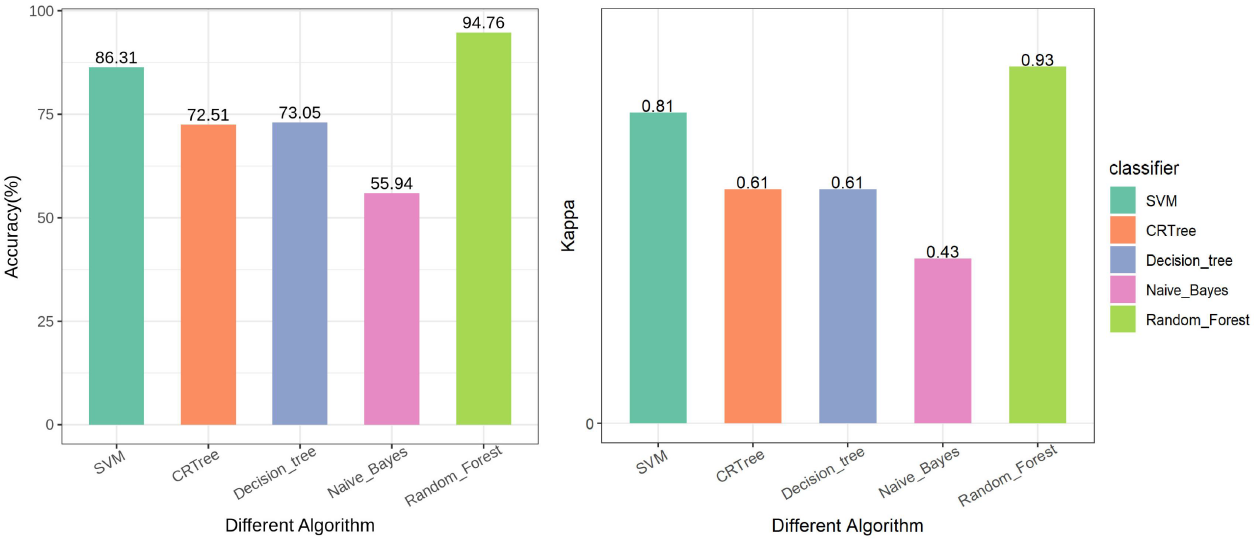
Accuracy rate and kappa coefficient of five machine learning algorithms on the testing set Note: SVM: Support vector machine algorithm, CTree: conditional inference tree algorithm, Decision_tree: decision tree algorithm, Naive_Bayes: Naive Bayes algorithm, Random_Forest: random forest algorithm.

**Figure 3.**
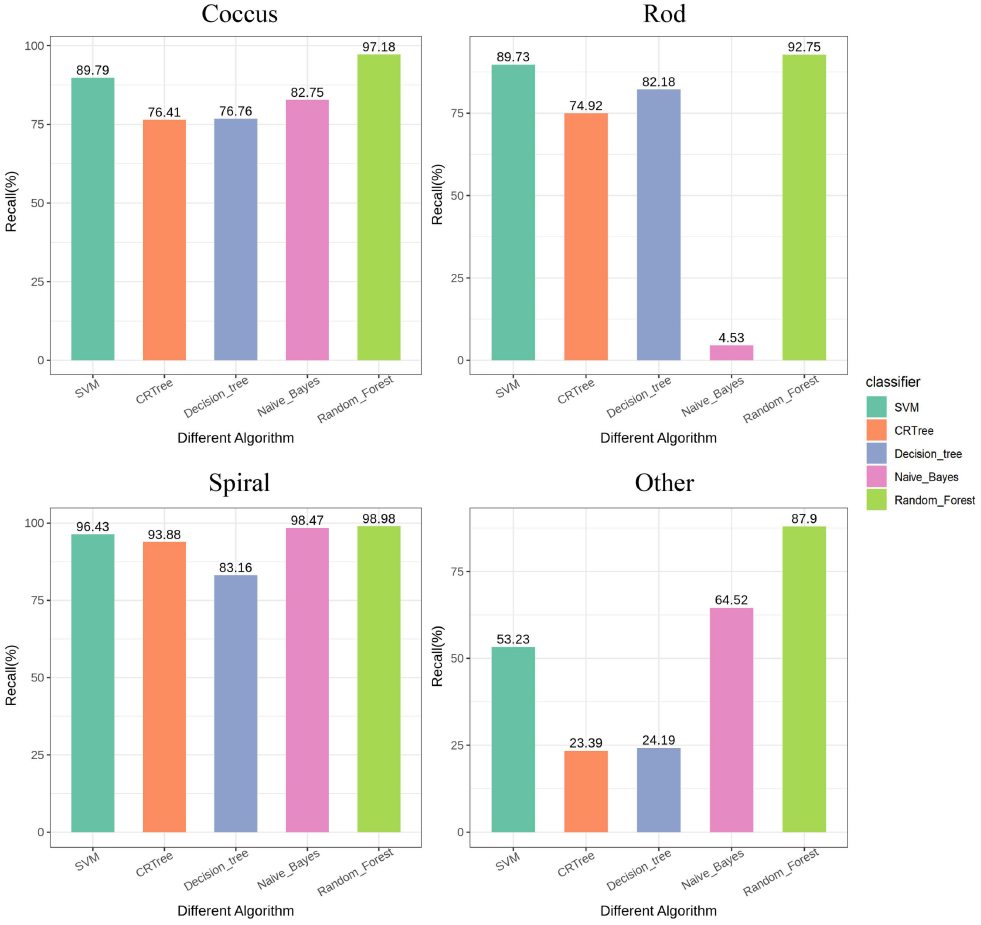
Recall rate of five machine learning algorithms on the testing set of different shape bacteria Note: SVM: Support vector machine algorithm, CTree: conditional inference tree algorithm, Decision_tree: decision tree algorithm, Naive_Bayes: Naive Bayes algorithm, Random_Forest: random forest algorithm.

### Methods for mining key genes

In our study, we focused on the rod shape as the trait of interest and utilized protein structure domains as the classification feature values in the previous machine learning predictions. To evaluate the importance of these classification features and identify potential key genes regulating rod shape, we employed the “importance” function in the random forest algorithm. To assess the importance of the classification features, we utilized the rfcv function in the randomForest program package to perform a five-fold cross-validation of the training model. This analysis generated a model accuracy curve (Figure 4), where the horizontal axis represents the number of feature values (specifically, protein structural domains in this study), and the vertical axis represents the error rate. Observing the curve, we noticed a significant decrease in the error rate as the number of feature values increased, particularly for the first 10 or so feature values. However, the curve tends to level off after surpassing 10 feature values, indicating that further increasing the number of feature values has little impact on the model accuracy. This suggests that the subsequent addition of feature values holds relatively low importance. Based on this analysis, we selected 10 feature values for this study as they were deemed most relevant and informative for predicting bacterial rod shape.

**Figure 4.**
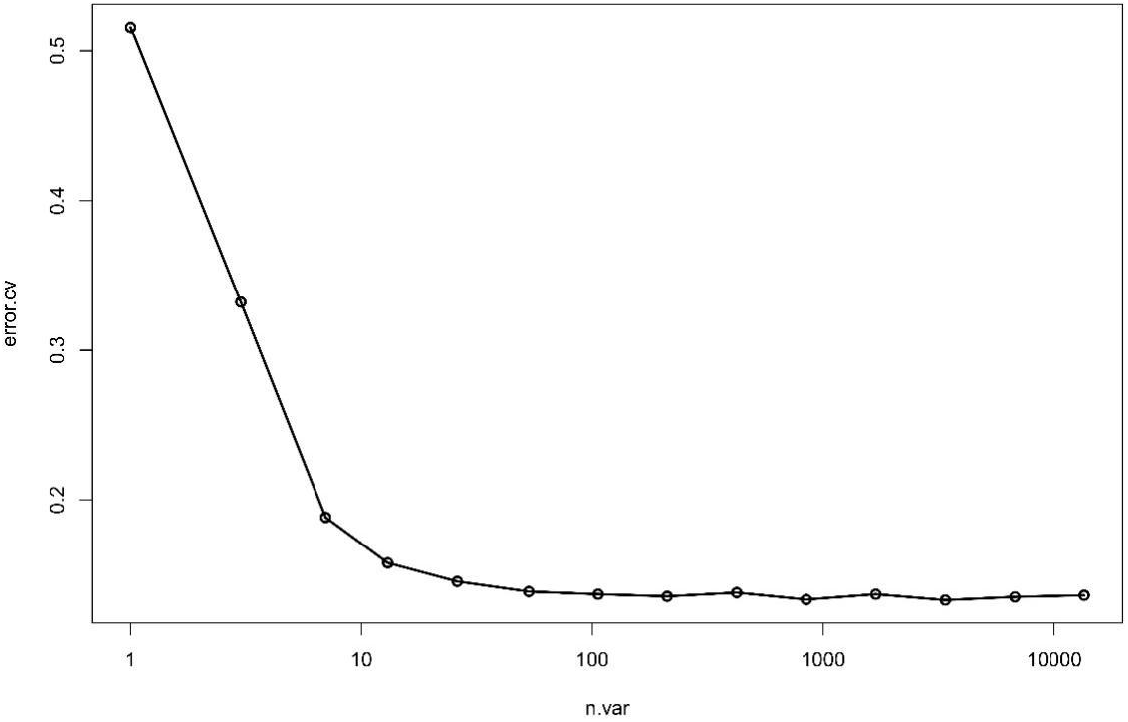
Model accuracy curve

The feature classification table of the optimal model is sorted in descending order according to the morphology “rod”. The top ten ranked structural domains are selected as the key determinant domains for bacterial rod shape (Table 2). The reference genome, protein, CDS, and other files of *E. coli* BL21 (DE3) strain (GenBank: GCA_013135395.1) are downloaded from NCBI. A self-developed program is used to compare the corresponding genes in *E. coli* BL21 (DE3) as key candidate genes. After screening, two of the top ten ranked structural domains, FlaA and FlgD, do not exist in the *E. coli* genome. Therefore, the remaining eight domains, pal, yicC, mreB, tolQ, ftsX, amiC, yddB, and rpoZ (Table 3), are selected as key candidate genes for subsequent gene function verification.

**Table 2.** Candidate domain of regulating bacterial length.

| Domain | Coccus | Rod↓ | Spiral | Other | MeanDecrease Accuracy | MeanDecrease Gini |
| --- | --- | --- | --- | --- | --- | --- |
| OmpA | 0.005285 | 0.002413 | 0.005321 | 0.00515 | 0.004262 | 6.238621 |
| YicC_N | 0.001397 | 0.002047 | 0.001862 | 0.003572 | 0.002016 | 1.886309 |
| FlaA | 0.000973 | 0.001771 | 0.011123 | 0.001188 | 0.003407 | 5.90406 |
| MreB_Mbl | 0.007044 | 0.001759 | 0.00416 | 0.003285 | 0.004038 | 4.691432 |
| MotA_Exb<br>B | 0.007018 | 0.001662 | 0.004844 | 0.004291 | 0.004293 | 5.899423 |
| FtsX_ECD | 0.000809 | 0.001579 | 0.004971 | 0.003328 | 0.002283 | 2.331755 |
| Amidase_3 | 0.000697 | 0.001513 | 0.000851 | 0.001527 | 0.00113 | 1.499186 |
| Plug | 0.001401 | 0.001462 | 0.001479 | 0.0014 | 0.001435 | 1.521337 |
| FlgD | 0.003965 | 0.001427 | 0.005393 | 0.004115 | 0.003401 | 3.466897 |
| RNA_pol_<br>Rpb6 | 0.000605 | 0.001347 | 0.005913 | 0.001938 | 0.002155 | 4.183334 |

**Table 3.** Candidate gene of regulating bacterial length.

| Domain | Genes | Protein id |
| --- | --- | --- |
| OmpA | *pal* | WP_001295306.1 |
| YicC_N | *yicC* | WP_000621336.1 |
| FlaA | - | - |
| MreB_Mbl | *mreB* | WP_000913396.1 |
| MotA_ExbB | *tolQ* | WP_001307062.1 |
| FtsX_ECD | *ftsX* | WP_001042003.1 |
| Amidase_3 | *amiC* | WP_000016907.1 |
| Plug | *yddB* | WP_000832450.1 |
| FlgD | - | - |
| RNA_pol_Rpb6 | *rpoZ* | WP_000135058.1 |

Note: -, It means it's not present in *E. coli*

### Morphological changes in *E. coli* after gene knockout

After validating the knockout genes through sequencing (Supplementary Figure 2), the knockout bacteria were subjected to scanning electron microscopy for observation. Among all the knockout strains, only *Δpal* and *ΔmreB* showed significant changes in the length or morphology of the bacterial cells compared to the original cells (Figure 5). The original *E. coli* cells exhibited long or short rod shapes with rounded ends. In contrast, the knockout strain lacking the *MreB* gene displayed a nearly spherical shape with a significantly shorter length and no notable change in diameter (Figure 6). The *E. coli* strain lacking the *Pal* gene became shorter in length, larger in diameter, and irregular in shape, resembling a protoplasmic state with no cell wall (Figure 6).

**Figure 5.**
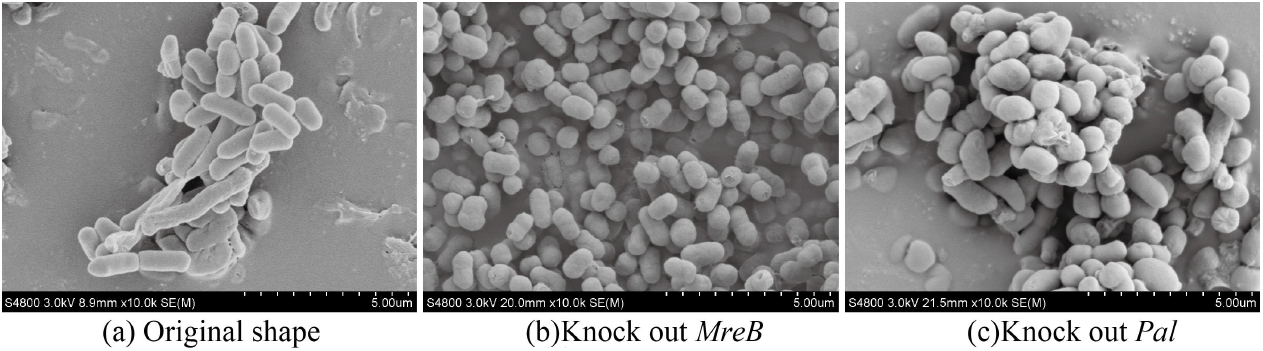
Morphology of *E. coli* after knockdown of *pal* and *mreB* genes, respectively Note:The magnification is 10,000 times, the length of the ruler is ten squares in total

**Figure 6.**
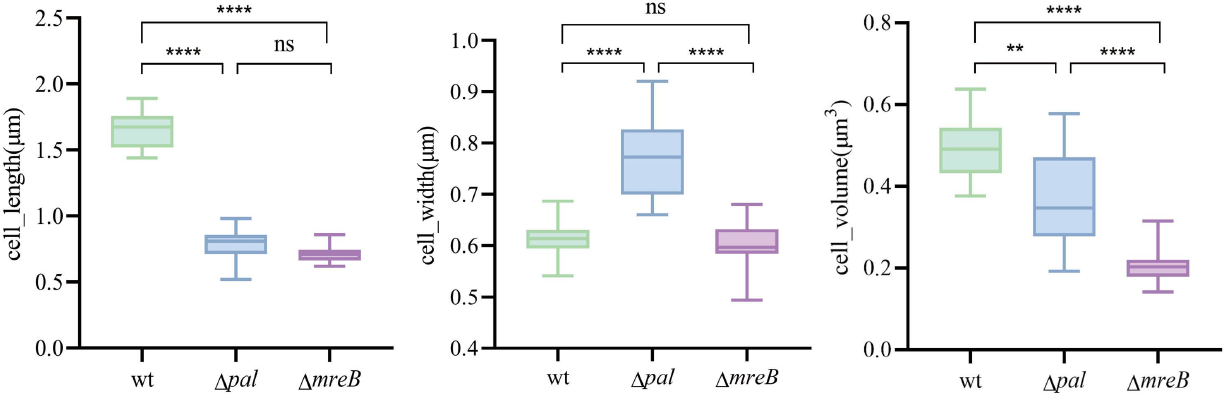
Bacterial cell size comparison Note: The F test showed that ns was not significant (p>0.05);** means significant (p<0.01); **** means extremely significant (p<0.0001)

Regarding the knockout strains of the remaining six genes, there were no significant changes observed in their length. However, some strains exhibited alterations in surface morphology, such as folds and depressions (Supplementary Figure 3). Knocking out the *yicC, tolQ, amiC, yddB*, and *rpoZ* genes resulted in the formation of multiple folds and depressions on the surface of *E. coli*. These five genes appeared to affect the outer wall morphology of the bacteria. On the other hand, the *ΔftsX* knockout strain maintained a smooth and rounded rod shape, showing no significant difference compared to the original strain (Supplementary Figure 3).

## Discussion

### Identification of key genes using multi-species genomes

Genes play a pivotal role in expressing life processes and characteristics, and their significance lies in unraveling the nature of life, exploring organismal development and evolution, and understanding the mechanisms behind disease occurrence. However, current methods for functional gene research primarily focus on analyzing individual species, and there is a need for new identification methods to discover functional genes that have significant effects on phenotypic traits. In contrast to traditional approaches involving mutant creation and population genetics, the novel technique established in this paper harnesses the vast amount of published genomic information directly, with lower cost and shorter time requirements.

In different species, even genes with the same function can possess highly distinct gene sequences, i.e. orthologous genes[30]. Identifying different sequences with the same function across diverse species is the key to the success of this new method. In this study, we employed protein structural domains as a means to distinguish genes with similar functions in different species[31], leading to a breakthrough in predicting bacterial shape. While some studies have utilized similar methods to establish correlations between genomes or proteins and traits, there is currently no definitive evidence confirming a direct relationship between the two[32].

In the initial machine learning prediction model, we utilized the protein structural domain as a classification feature value. By employing the “importance” feature in the random forest algorithm[33-35], we assessed the significance of each classification feature. A higher importance ranking indicated a greater impact of the feature in the classification algorithm, specifically in determining the shape of bacteria. Based on this importance index, we hypothesized that these genes could be essential combinations of genes related to a specific trait. To validate this conjecture, we conducted experimental validation by individually knocking out the first eight representative genes. Our observations revealed significant changes in the length or morphology of *E. coli* after knocking out the *Pal* and *MreB* genes. Specifically, the knockout of *MreB* transformed the bacteria from rod-shaped to spherical, which is consistent with findings from previous studies[9]. Additionally, the knockout of *YicC, TolQ, AmiC, YddB*, and *RpoZ* genes resulted in the formation of folds and depressions on the surface of *E. coli*. These observations suggest that the key genes identified in this study, through rapid mining of key functional genes, do indeed play a role in determining the rod shape of bacteria. Furthermore, the analytical method established in this paper, which involves mining key genes regulating common features using genomic features from multiple species, appears to be a feasible approach.

### Optimization of the model

The relative proportions of different bacterial types have a significant impact on the prediction accuracy. Initially, we utilized all 4,847 bacterial species with available genomic and trait information for training and testing the model. However, the results were unsatisfactory, as indicated by poor Kappa coefficients. Specifically, both conditional inference tree algorithm and Naive Bayes algorithm yielded Kappa coefficients of 0, indicating inaccurate predictions. One potential explanation for this outcome is the high prevalence of bacilli in the bacterial dataset. To address this issue, we attempted to rebalance the dataset by reducing the number of bacilli and increasing the number of cocci and spirochetes. By doing so, we aimed to create a more representative and diverse dataset that could potentially enhance the accuracy of our predictions.

The dataset was refined by filtering bacilli to 1,380 representative species from the initial 3,991 bacilli species. Additionally, Bergey’s manual of Determinative Bacteriology [36], was consulted to manually add 251 species of cocci and 160 species of spirochetes. The resulting training set consisted of 534 cocci species, 1,035 bacilli species, 178 spirochetes species, and 239 other bacterial species. The test set comprised 177 cocci species, 345 bacilli species, 60 spirochetes species, and 79 other bacterial species. Therefore, the dataset consisted of 2,647 bacteria species, with 1,986 species in the training set and 661 species in the test set. Following these modifications, the optimized results showed improvements compared to the initial modeling outcomes. Among the five algorithms used, the support vector machine and random forest algorithm demonstrated better performance in the training set, achieving prediction accuracies of 85.60% and 76.03%, respectively. However, when predicting the test set, the support vector machine performed less effectively than the random forest algorithm.

By manually adjusting the number of strains for cocci, bacilli, and spirochetes and carefully examining the phenotypes, the training set was further refined. The revised training set consisted of 854 cocci species, 999 bacilli species, 589 spirochetes species, and 373 other bacterial species. The test set included 284 cocci species, 331 bacilli species, 196 spirochetes species, and 124 other bacterial species. The classification results from the five algorithms improved, and two algorithms achieved Kappa coefficients of 0.88 and 0.83 in the training set, namely the support vector machine and random forest algorithms, indicating a nearly perfect agreement between the predicted and true values. In terms of accuracy and recall on the training set, the support vector machine exhibiting the highest performance, followed by the random forest algorithm. However, when predicting the test set, the support vector machine’s performance declined, and the random forest algorithm emerged as the top performer, achieving an accuracy of 94.76% and recall rates of 97.18%, 92.75%, 92.75% for cocci, bacilli, spirochetes, and other bacteria, respectively. The Kappa coefficient reached 0.93, indicating a strong prediction capability. Obtaining high prediction accuracy is likely the key to identifying key genes.

### Key genes regulating bacterial rod shape

The *E. coli* MreB protein which is one of the main components of the cell cytoskeleton and plays a crucial role in cell vitality and the formation of rod-shaped cell morphology[9]. The MreB protein forms helical filaments underneath the cell membrane in *E. coli* and controls cell wall growth and maintenance of cell shape[37, 38]. Mutations in this gene can lead to loss of cell wall symmetry and the appearance of bent or abnormally twisted cell morphologies[39]. Therefore, in our study, the complete knockout of the *MreB* gene affected cell wall synthesis and assembly, causing *E. coli* to lose the cell cytoskeleton and resulting in cell death (Data not show). Some partially knocked out bacteria were able to survive, possibly due to the retention of partial cell cytoskeleton functionality, which allows them to maintain cellular activity. However, this also affects their ability to maintain their morphology, resulting in a shorter rod-shaped or nearly spherical morphology in *ΔmreB* mutants.

The *E. coli* Pal protein which is associated with peptidoglycan[40]. The Pal protein interacts with the BamA protein, participating in the folding and assembly processes of outer membrane proteins and aiding in the maintenance of outer membrane integrity[41, 42]. Deletion of the *Pal* gene leads to outer membrane abnormalities and increased sensitivity to antibiotic drugs[40]. The outer membrane is one of the main determining factors of bacterial morphology[43]. Therefore, in this study, the *ΔPal* mutant may have altered the outer membrane of *E. coli*, resulting in changes in its shape, such as becoming nearly spherical or irregular. The knockout of the *YicC, TolQ, AmiC, YddB*, and *RpoZ* genes did not cause significant changes in the length or shape of *E. coli*, but it did result in the formation of many folds and depressions on the bacterial surface. This suggests that these genes also have some impact on the bacterial phenotype, but their effects may require two or several of them to act in combination.

The deletion of the *Pal* and *MreB* genes not only altered the cell morphology of *E. coli* but also affected the growth rate and cell size of the bacteria. It has been reported that the MreB protein localizes densely in the cylindrical part of *E. coli* cells, rather than in the polar regions[44]. The MreB protein affects the bidirectional elongation of the cell by regulating peptidoglycan synthesis, thereby influencing polar growth of the cell and exerting an impact on the polarity of cell growth[12, 45, 46]. The Pal lipoprotein plays a major role in the integrity of the outer membrane in *E. coli*[40]. Therefore, after losing the *Pal* gene, the growth rate of *E. coli* significantly slows down, and the changes in length and diameter are likely due to the incomplete outer membrane caused by the loss of the *Pal* gene, resulting in a protoplasmic state and the formation of irregular “spherical” *E. coli* cells.

In addition to validating the two genes, *Pal* and *MreB*, which influence rod-shaped bacterial morphology in this study, other genes known to affect bacterial morphology, such as *RodZ* and *PBP*[47, 48], were not directly obtained. One possible reason is that the shapes obtained from online sources or through book references and extrapolation may still contain errors. In the early stages, a large number of genomes were obtained through second-generation sequencing, and there may still be room for improvement in the accuracy and completeness of the assembled genomes. These factors may have contributed to the lack of precision in the model established in this study, which could directly affect the accurate identification of key genes. Although we were unable to obtain the complete set of genes encoding rod shape, the simultaneous identification of two key rod-shaped genes represents a significant technological advancement. With the progress of third-generation sequencing technologies and the completion of more comprehensive and extensive genomes, we believe that the methods we have developed will assist more researchers in rapidly and accurately identifying key genes.

### Materials and Methods Bacterial genomes

Bacterial reference genome data were downloaded from the NCBI (https://www.ncbi.nlm.nih.gov/) database to a local laboratory server for a total of approximately 224,000 bacterial reference genomes (October 2022).

### Construction of eigenvalues

Protein structural domain resolution of genomic data was performed using the software package pfam_scan[49], Pfam-A database version is 33.0. To analyze the resolved protein domains of bacteria, we concatenated the protein domains obtained from each bacterium’s genes. The concatenated protein domains were used as row names in the analysis. We then counted the occurrence of each concatenated domain and integrated the counts to construct a frequency matrix of protein domains for each bacterium. This process allowed us to create a dataset containing characteristic values for multiple bacterial protein domains.

### Constructing a feature classification table

From the bacterial information query website, BacDive (https://bacdive.dsmz.de/), we obtained information on the known shapes of bacteria. As of June 2022, there are a total of 4,847 bacterial species with shape information. Among them, 3,991 species are rod-shaped bacteria, followed by 460 species of spherical bacteria, and 79 species of spiral bacteria. There are also 317 species with other shapes such as star-shaped, filamentous, brick-shaped, pleomorphic, and ring-shaped. To facilitate the construction of the model, we categorized the bacterial shapes into four classes based on the classical classification method: cocci for spherical bacteria, rods for rod-shaped bacteria, spiral for spiral bacteria, and others for the remaining shapes.

### Machine learning model construction and optimization

The three essential elements for building a machine learning model are feature values, grouping lists, and algorithms. We obtained a feature value dataset by intersecting the bacteria with known shape information and the frequency matrix of protein domains for the bacteria based on their names. Among the 4,847 bacterial species, we divided them into training and testing sets in a 3:1 ratio, resulting in the grouping list. To select the best algorithm, we compared five different algorithms: decision trees[50], random forests[33], support vector machines[51], Conditional Random trees[52], and naive Bayes[53].

In the field of machine learning, it is common to use evaluation metrics to assess the performance of classification models. Three commonly used evaluation metrics are accuracy, recall, and Kappa coefficient[54, 55]. These metrics are calculated and analyzed based on the confusion matrix generated from predictions. Based on these metrics, continuous adjustment and optimization of the model are needed to achieve better classification performance and accuracy. In this study, the main adjustment is the proportion of different types of bacteria.

### Identifying key candidate genes

In the machine learning prediction results, we used protein domains as classification features. The “importance” function in the random forest algorithm can be used to assess the importance of classification features. The higher the ranking of importance, the more significant the feature is in the classification algorithm[35]. This indicates that the protein domain has a greater influence on determining bacterial shape. It forms the theoretical basis for this study to explore key shared genes using a large amount of genomic information from different species.

*E. coli* BL21 (DE3) was selected as the original strain for subsequent knockout experiments. The top ten ranked protein domains were chosen as key determinants of bacterial rod shape. Homology alignment and the relationship with Pfam protein domains were used to identify corresponding genes as key candidate genes.

### Gene function validation

Gene editing of candidate genes in *E. coli* BL21 (DE3) strain was performed using CRISPR-cpf1 technology. Scanning electron microscopy (SEM) was used to observe changes in the shape of the knockout strains, and growth curves were determined for the strains that showed significant shape changes before and after knockout.

A dual-plasmid gene editing chemical transformation system pEcCpf1/pcrEG based on CRISPR/Cpf1 in *E. coli* was used in this study[56]. Throughout the process, the final concentrations of antibiotics used were kanamycin at 50 µg/ml and spectinomycin at 100 µg/ml, and the cultivation temperature was maintained at 37 °C. The morphology of *E. coli* before and after gene knockout was observed using scanning electron microscopy.

Electron microscopy images of the experimental group and control group at a magnification of 10,000x were selected. The measurement tool in image processing software was used to set the measurement scale as “40 dpi = 0.2 µm”. Sixteen independent measurements of length (l) and diameter (d) were taken for each strain, and the values were recorded. The volume was calculated using formula 3-1.

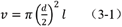

